# Abnormal Vaginal Microbiota Associated with miRNA Targeting the HIV-Host Interactome

**DOI:** 10.1101/2025.07.18.665639

**Authors:** Raina N. Fichorova, Paula F. T. Cezar-de-Mello, Jonathan M. Dreyfuss, Hidemi Yamamoto, Pai-Lien Chen, Pan Hui, Xiaoming Gao, Charles Morrison, Robert Barbieri, Gustavo F. Doncel

## Abstract

Understanding the molecular mechanisms underlying the ability of vaginal dysbiosis to alter the mucosal barrier to HIV acquisition is an essential step toward prevention. We hypothesized that micro(mi)-RNAs dysregulated by vaginal pathobiont bacteria epigenetically control host pathways exploited by the virus. The impact of these endogenous non-coding short RNAs on the anti-viral mucosal barrier function in the female reproductive tract is largely unknown. This study utilized cervicovaginal specimens collected during the luteal and follicular phase of the menstrual cycle along with data on age, race, ethnicity, education, and body mass index from 141 healthy reproductive-age women confirmed negative for sexually transmitted infections. Vaginal microbiota was classified by Nugent scoring. Shot-gun vaginal microbiome sequencing and metagenome taxonomic classification was performed on a subset of 21 women. Levels of miRNAs in exosomes isolated from cervicovaginal secretions were quantified using the EdgeSeq-NextGen global transcriptome platform. Differential expression (DE) was determined using R. Epigenetic target prediction was performed using MirTarBase. MiRNA profiles varied by both Nugent score categories (0-3 scores = normal, 4-6 = intermediate, and 7-10 = bacterial vaginosis, BV) and by metagenome classification. Higher microbiome diversity was associated with higher number of significantly dysregulated miRNAs (588 in BV compared to Nugent 0-3 versus 42 in Nugent 4-6 compared to Nugent 0-3, false discovery rate FDR<0.01) affecting over 400 experimentally validated genes targeted for post-transcriptional regulation. The miRNAs dysregulated by *G. vaginalis*-dominated compared to *L. crispatus*-dominated metagenomes included 24 DE miRNAs (92% overlap with BV by Nugent score) and 112 validated target genes. BV-dysregulated miRNA mediated the immunosuppressive effects of BV on cytokine levels previously associated with HIV acquisition risk. The gene ontology predictions based on BV-dysregulated miRNAs identified enrichment for 445 downregulated and 50 upregulated genes previously validated as part of the HIV-host interactome. miRNAs mediation revealed a mechanism of suppressed immunity by BV predictive of HIV risk. In conclusion, miRNAs dysregulated by vaginal dysbiosis may facilitate immune imbalance and cellular pathways associated with HIV risk.

## 1 Introduction

Disturbances in the vaginal microbiota composition classified by either Nugent scores or microbiome community types (1) has been clinically associated and experimentally validated as a significant cause of breaking the antiviral mucosal host defense against HIV (2). Therefore, understanding the molecular mechanisms underlying the role of the vaginal microbiota in HIV acquisition risk is an essential step toward safer and more effective HIV prevention. We hypothesized that micro(mi)-RNAs regulated by the resident microbiota, a mechanism understudied to date, can interfere with host pathways exploited by the virus. Micro-(mi)RNAs are endogenous short non-coding RNA molecules that exert post-transcriptional epigenetic regulation by blocking or limiting mRNA translation into protein. They are selectively packaged within small extracellular vesicles (EV) (exosomes) or form stable complexes with proteins to mediate signaling between distant anatomic sites(3). Because of their stability in the peripheral circulation and their established epigenetic function they are attractive therapeutic and diagnostic targets. Recently, miRNAs and exosomes were proposed to be important players in HIV pathogenesis(4) and in bacterial infection (5); however, comprehensive analysis of the cervicovaginal miRNA EV cargo in healthy reproductive age women has not been reported to date. This study pioneers the filed by applying global transcriptome analysis of EV isolated from cervicovaginal secretions of healthy non-pregnant reproductive-age women to identify miRNAs differentially dysregulated by vaginal microbiota classifications. It hypothesized that vaginal dysbiosis will be associated with a) unique EV miRNA profiles, and b) miRNA-mediated enrichment for cytokine markers of HIV acquisition risk as well as for pathways of the global host-HIV interactome defined as genes validated to take part in cellular events of HIV uptake and pathogenesis.

## 2 Materials and Methods

### 2.1 Study population and ethics statement

The study was performed in agreement with the Declaration of Helsinki for ethical conduct of human subjects’ research. All participants signed informed consent and the study was approved by the Institutional Review Boards of Eastern Virginia Medical School and Brigham and Women’s Hospital.

The study performed global miRNA transcriptome analysis on 288 cervicovaginal specimens from 141 healthy reproductive-age women collected from four clinical trials conducted by the Contraceptive Research and Development Program (CONRAD) at Eastern Virginia Medical School. The clinical and demographic characteristics of the study population are presented in **Table 1**.

**Table 1.** Demographic characteristics of 140 study participants (one participant is missing baseline information)

|  | By Race |  |  |  | By Ethnicity |  |  |  |
| --- | --- | --- | --- | --- | --- | --- | --- | --- |
|  | Black | White | Mixed/<br>Other | Total | Hispanic or<br>Latina | Not<br>Hispanic or<br>Latina | Not<br>Reported | Total |
| Participants | 62 (44.3%) | 49 (35.0%) | 29 (20.7%) | 140 | 29 (20.7%) | 77 (55.0%) | 34 (24.3%) | 140 |
| Age |  |  |  |  |  |  |  |  |
| Mean±SD<br>(range) | 34.2±7.9<br>(18.0 -<br>51.0) | 31.3±7.8<br>(20.0 -<br>49.0) | 31.7±7.3<br>(19.0 -<br>49.0) | 32.7±7.8<br>(18.0 -<br>51.0) | 30.6±7.0<br>(18.0 -<br>46.0) | 33.1±8.4<br>(19.0 -<br>51.0) | 33.5±7.0<br>(19.0 -<br>49.0) | 32.7±7.8<br>(18.0 -<br>51.0) |
| Age < 25 | 9 (14.5%) | 12 (24.5%) | 5 (17.2%) | 26 (18.6%) | 6 (20.7%) | 17 (22.1%) | 3 (8.8%) | 26 (18.6%) |
| Age >=25 | 53 (85.5%) | 37 (75.5%) | 24 (82.8%) | 114<br>(81.4%) | 23 (79.3%) | 60 (77.9%) | 31 (91.2%) | 114<br>(81.4%) |
| Education years |  |  |  |  |  |  |  |  |
| <= 12 | 18 (31.0%) | 10 (23.8%) | 18 (64.3%) | 46 (35.9%) | 19 (67.9%) | 20 (30.3%) | 7 (20.6%) | 46 (35.9%) |
| > 12 | 40 (69.0%) | 32 (76.2%) | 10 (35.7%) | 82 (64.1%) | 9 (32.1%) | 46 (69.7%) | 27 (79.4%) | 82 (64.1%) |
| BMI categories |  |  |  |  |  |  |  |  |
| Underweight | 1 (1.6%) | 1 (2.0%) | 1 (3.4%) | 3 (2.1%) | 1 (3.4%) | 1 (1.3%) | 1 (2.9%) | 3 (2.1%) |
| Healthy Weight | 11 (17.7%) | 24 (49.0%) | 8 (27.6%) | 43 (30.7%) | 8 (27.6%) | 25 (32.5%) | 10 (29.4%) | 43 (30.7%) |
| Overweight | 16 (25.8%) | 15 (30.6%) | 9 (31.0%) | 40 (28.6%) | 9 (31.0%) | 22 (28.6%) | 9 (26.5%) | 40 (28.6%) |
| Obese | 34 (54.8%) | 9 (18.4%) | 11 (37.9%) | 54 (38.6%) | 11 (37.9%) | 29 (37.7%) | 14 (41.2%) | 54 (38.6%) |

All four trials enrolled reproductive age women (18-49 years old) who gave consent for the future research use of the banked specimens. Data were consistently collected between the trials on age, race, and body mass index (**Table 1**). Details on each clinical protocol can be found in published reports (6-9) and at <u>ClinicalTrials.gov</u>. Participants were asked to refrain from vaginal intercourse 48 hours before all sampling visits. All subjects were confirmed negative for sexually transmitted infections at the time of sampling. Negative tests included a pap smear (Thin Prep, Hologic, Bedford, MA), *Neisseria gonorrhoeae* and *Chlamydia trachomatis* (Gen-Probe, Aptima, San Diego, CA). Point-of-care testing was performed to exclude pregnancy, recent vaginal semen exposure (ABAcard, Abacus Diagnostics, West Hills, CA), HIV-1 infection (Oraquick Advance, Orasure Technologies, Bethlehem, PA), and *Trichomonas vaginalis* infection (OSOM, Genzyme, Cambridge, MA). The presence of active genital herpes was ruled out by clinical exam. All studies were conducted before the emergence of SARS-Cov2.

Baseline visits (prior to any intervention) were contributed by the following trials: “A Phase I Safety Study of the Caya Diaphragm Used with ContraGel” (protocol # A13-127) (9), “Pharmacokinetic and Pharmacodynamic Study of Tenofovir 1% Gel Using the BAT 24 Regimen Versus Daily and Pericoital Dosing” (protocol # A10-113) (10), “Assessing the Effect of Contraception and the Menstrual Cycle on Pharmacokinetics, Pharmacodynamics, and Vaginal Safety in Tenofovir Vaginal Gel Users” (protocol # A10-114) (7), and “Biomarkers of Bacterial Vaginosis” (protocol # A13-115) (8). The A13-115-study enrolled and treated with metronidazole 33 women diagnosed with BV by Nugent score and followed them testing for cure or recurrence at one week and one month after treatment (8). In addition to specimens from the baseline visit, this study contributed for miRNA analysis the paired samples collected at the follow up visits.

### 2.2. Biospecimen processing and EV isolation

While different methods were used for collection of cervicovaginal secretions, the isolation of extracellular vesicles provided a uniform vehicle for quantitation of relative miRNA expression levels within each biospecimen. Study #A10-114 (6) obtained cervicovaginal fluids by direct aspirate from the posterior vaginal fornix with a 2.5-mL vaginal fluid aspirator (CarTika Medical, Maple Grove, MN) and immediately transferred to a cryovial and stored at −80°C. Studies #A10-113 and #A13-115 used cervicovaginal lavage (CVL) performed with 10 cm^3^ of normal saline (8). Within 30 min of collection, the CVL was centrifuged at 4°C for 10 min at 500×g and supernatants stored at −80°C. Study #A13-127 collected cervical and vaginal swabs which were snap frozen in liquid nitrogen prior to elution in Fichorova’s lab in 1 ml PBS, aliquoted and frozen at -80°C. CVLs and swab elution supernatants were mixed in a 2:1 v/v ratio with the Total Exosome Isolation Reagent (Invitrogen, Carlsbad, CA) and incubated for 16 h at 4°C followed by centrifugation using a fixed angle rotor at 10,000 x g for 60 minutes at 4°C. ZetaView (Particle Metrix, Meerbusch, Germany) was used to assess size and concentration of EVs and transmission electron microscopy with immunogold labelling of CD63 was used to confirm the presence of exosomes in the EVs samples as previously described (11). For miRNAs expression profiling, supernatants were removed and EV pellets were resuspended in 35 µl of HTG lysis buffer to further miRNA transcriptome assessment.

### 2.3 Vaginal microbiota classification by Nugent score and shotgun sequencing

Vaginal microbiota was classified by Nugent scoring (12) at each study visit. Microbiome metanalysis was performed on all baseline specimens collected from clinical trial #A13-127. DNA was isolated from swab elution cell pellets using ZYMObiomics Microprep Kit, according to the manufacturer’s protocol. Isolated DNA was quantified by Qubit. Fragment libraries were generated for the DNA using the Thermo Fisher IonXpress Plus Fragment Library kit according to the manufacturer’s instructions. The libraries were then sequenced on an Ion S5XL sequencer to generate 200bp sequences. Unassembled sequencing reads were directly analyzed by CosmosID bioinformatics platform (CosmosID Inc., Rockville, MD) described elsewhere (13-16). Briefly, the system utilizes curated genome databases and a high-performance data-mining algorithm that rapidly disambiguates hundreds of millions of metagenomic sequence reads into the discrete microorganisms engendering the particular sequences.

### 2.4. Global miRNA transcriptome quantification and targets annotation

Levels of miRNAs were quantified in extracellular vesicles isolated from the cervicovaginal secretions using the High Throughput Genomics (HTG) EdgeSeq global transcriptome platform (HTG Molecular Diagnostics Inc, Tucson, AZ). The library was constructed using automated high-fidelity cost-effective quantitative nuclease protection assay (qNPA) coupled with next generation sequencing (NGS). The resulting libraries were sequenced by NextSeq using Illumina. A plant gene (*Arabidopsis thaliana aintegumenta*, ANT, accession no. U41339) was included in each assay as a negative background control. Fifteen housekeeper genes were used for a robust control of gene expression (ACTB, ATP5F1, DDX5, EEF1G, GAPDH, NCL, OAZ1, PPIA, RPL38, RPL6, RPS7, SLC25A3, SOD1, TBP and YWHAZ). Each sequencing run included quality control samples with known levels of RNA expression.

After filtering out miRNAs that have at least 5 counts per million (CPM) in at least 5 samples, counts for 2801 miRNAs were normalized by the upper-quartile method to align all samples’ 0.95 percentile (17). Voom transformation (18) was applied to obtain normalized log_2_CPM. Differentially expressed (DE) miRNA between groups were identified using moderated t-tests from the R package Limma (19), which applies linear regression models. Adjusting for multiple measured covariates, such as age, BMI, race/ethnicity, and education was considered. Analysis with the R package variancePartition (20) showed that these covariates explain too little of the variation in microRNA expression to be worthwhile to include, and some had many missing values. Therefore, no adjustment was performed for any covariates. The analysis accounted for multiple hypotheses using the Benjamini-Hochberg false discovery rate (FDR) procedure with a significance cut-off of FDR 1%. The miRNA statistics for the full miRNAome is provided in **Supplemental Table 1** which contains the unweighted average logCPM for each comparison group, weighted p values and FDR from the moderated t test, log fold change and fold change of each comparison, and miRNA annotation based on miRbase v.20 and v.22. All miRNAs were annotated from miRBase v20 to miRBase v22 accession (21) using miRBaseConverter (v.1.22.0) (22). Sixteen miRNAs, considered *dead* in miRBase v22, were excluded from the target analysis. MiRNAs were then mapped to their targets in miRTarBase (23), focusing on those validated by a luciferase reporter assays to be most confident that the interactions were direct.

### 2.5. Cytokines – miRNA interactions

Levels of SLPI, IL-1RA, IL-8, IL-6 and RANTES were previously measured in the CVLs and cervical swab elutions from all 4 clinical trials as described (7-9). These mediators of inflammation and innate immunity were chosen based on published evidence for their association with BV and HIV risk and experimental evidence for causal relationship with BV associated bacteria (8, 24-30): lower SLPI associated with increased HIV risk and BV (8, 24, 30).

Cytokine levels were log2-transform to approximate normal distribution and ordered into tertiles into low, medium and high rank conditional on sample type e.g. vaginal swab or cervicovaginal lavage sample that was used for both cytokine measurement and EV isolation for miRNA analysis. Limma (v.3.54.0) was also used to test association of cytokines categories with continuous levels of miRNAs. Spearman correlation analysis was used to identify significant association of BV status with each cytokine.

High-throughput mediation analysis (Hitman, v.0.1.2.9010) (31) was performed to test the BV➔miRNA➔cytokine mediation for all miRNAs where exposure (E) is BV, the potential mediator (M) is miRNA and the outcome (Y) is the cytokine abundance. Hitman was not limited to miRNAs that are targeting the cytokine according to MirTarBase or miRNAs that are negatively associated with the cytokine. This approach allows exploration of both direct and indirect effects of the miRNA on the cytokine. The mediation analysis was limited to the cytokines that showed significant association with BV (P<0.05 and FDR <0.05). The statistics for all miRNAs are provided in **Supplemental Table 2.** For each cytokine, the columns are: EMY.chisq, which is the overall statistic for mediation where larger is stronger; EMY.p, which is the p-value for mediation; EMY.FDR, which is the FDR for mediation. We also have EM.z, which is the z-score for E–>M, not accounting for direction; EM.p, which is the p-value for E–>M, not accounting for direction; MY.z, which is the z-score for M–>Y, not accounting for direction; MY.p, which is the p-value for M–>Y, not accounting for direction.

We performed network analysis to map BV interactions with miRNAs directly regulating cytokine translation, selecting the candidate BV -> mir -> cytokine connections according to the following criteria: (a) the microRNA is affected by BV (Nugent 7-10 versus Nugent 0-3, FDR < 1%) and (b) the cytokine is the target of the miRNA based on TargetScan 7.2.

### 2.6. Pathway analysis with modular enrichment and overlap with the HIV interactome

Modular Enrichment Analysis (MEA) was applied to the validated targets of differentially expressed EV-microRNAs (DEmiRs) (FDR < 0.01) across Nugent score comparisons (BV vs. normal, intermediate vs. normal, BV vs. intermediate), alongside *G. vaginalis*-dominated compared to *L. crispatus*-dominated metagenomes. This analysis utilized ClueGo v2.5.9 (32) and CluePedia v1.5.9 (33) within Cytoscape v3.10.3. Non-redundant biological terms and their linked functionally associated networks (FAN) were identified from the Gene Ontology Immune System (GOIS). The gene sets were divided into Cluster 1 (upregulated) and Cluster 2 (downregulated). Enrichment significance was assessed using a two-sided hypergeometric test with Bonferroni step-down correction (p ≤ 0.05), and networks were visualized using the Organic Layout algorithm (yFiles Layout Algorithms, v1.1.4).

Additionally, MEA was performed on HIV interactome genes that overlap with validated targets of miRNAs dysregulated across all Nugent score comparisons and metagenome, as above. The HIV interactome genes were sourced from the NCBI database (34) and a previous report (35). This analysis followed the same methodology described above but incorporated both the GOIS and Kyoto Encyclopedia of Genes and Genomes (KEGG) databases.

## 3 Results

### 3.1. Differentially expressed EV-microRNA targets across Nugent score comparisons and per species-dominated metagenome

The cervicovaginal miRNA profiles varied by both Nugent score categories (0-3 scores – normal, 4-6 – intermediate, and 7-10 – bacterial vaginosis, BV) and by metagenome classification which revealed distinct microbiome clusters by Nugent score as illustrated by heatmaps and volcano plots in **Fig. 1** and **Fig. 2**, respectively (stats in **Supplemental Table 1**). **Table 2** lists the number of vaginal dysbiosis-regulated DE EV miRNAs, their targets experimentally validated by our most stringent validation criteria, direction of regulation and the number of validated targets overlapping with the HIV interactome. All validated targets of the DE EV-miRNAs are shown in the supplemental tables **Supplemental Table 3**.

**Figure 1.**
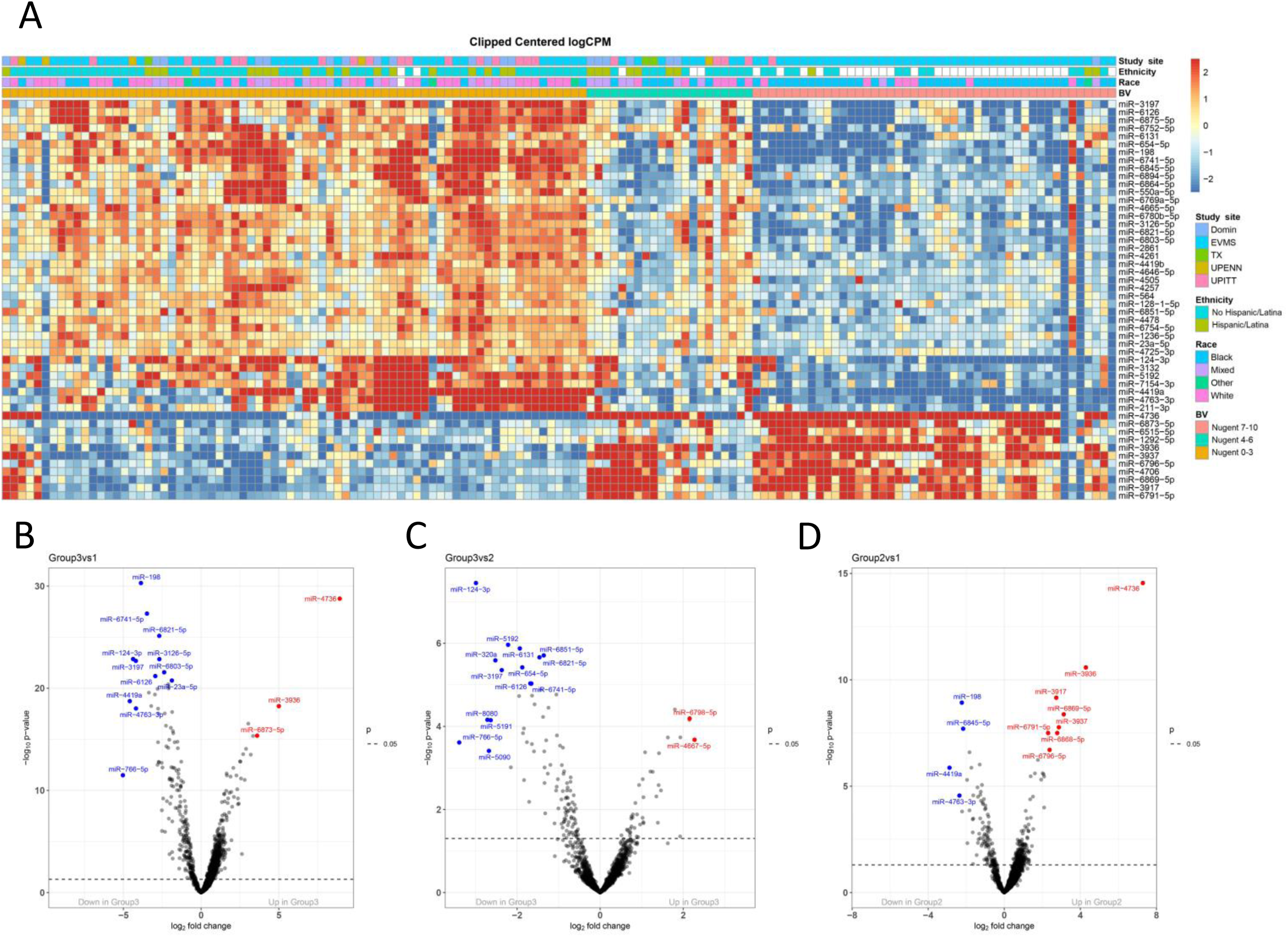
DE miRNAs distinguishing dysbiosis by Nugent score: (A). Heat map of top 50 DE miRNAs. (B-D) Volcano plots showing downregulated to the right and upregulated to the left miRNAs when comparing Nugent score 7-10 (group 3) to normal Nugent score 0-3 (group 1) (B) and to Nugent score 4-6 (group 2) (C), and when comparing Nugent score 4-6 to normal Nugent score 0-3 (D).

**Figure 2.**
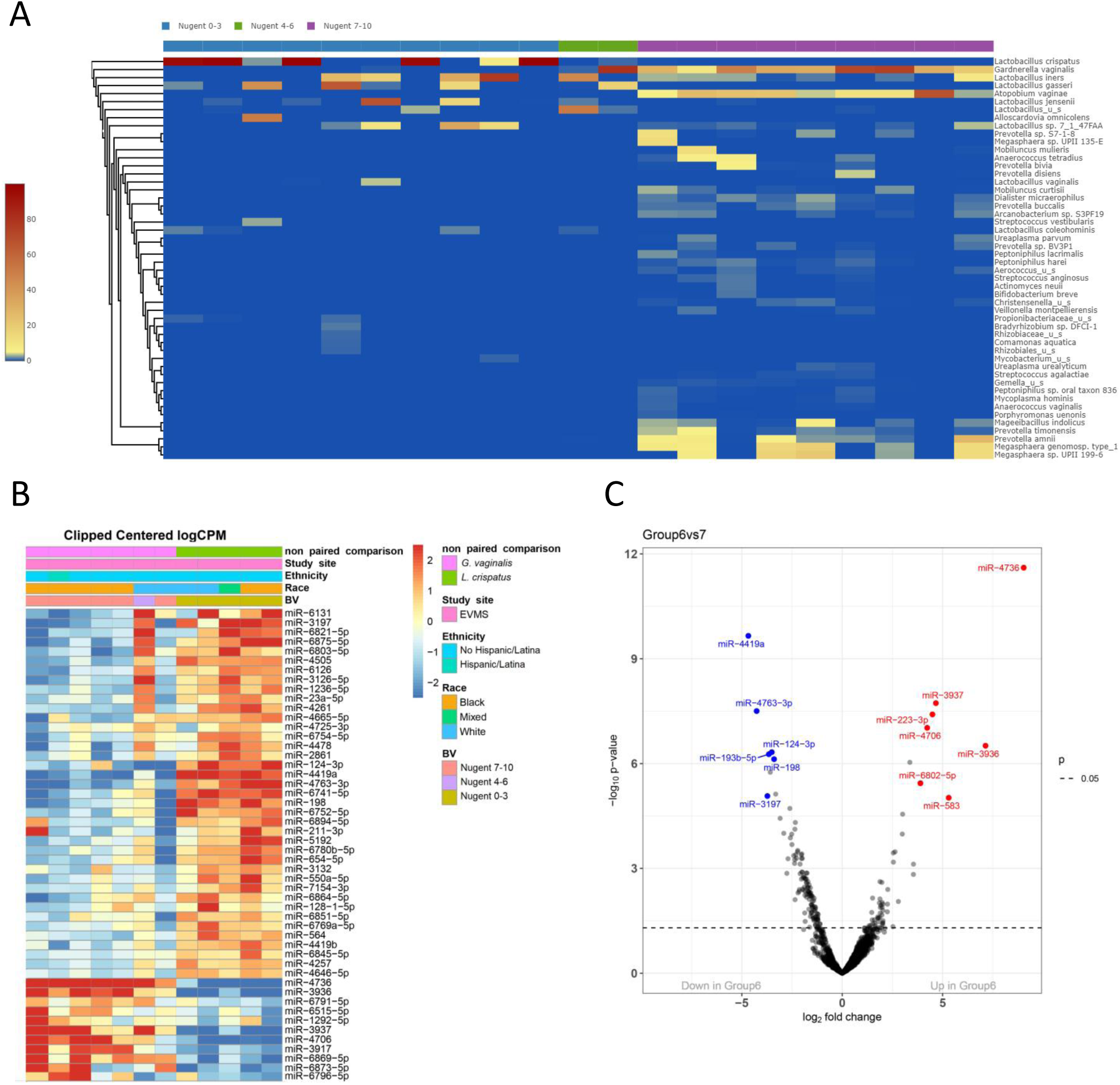
Microbiome metagenome shot gun sequencing-based clustering by Nugent Score: (A) Heatmap of differentially expressed miRNAs distinguishing dysbiosis Nugent 4-6 and 7-10 from normal Nugent 0-3. (B) Heat map of top 50 MiRNAs differentially regulated by the vaginal microbiome dominated by *G. vaginalis* (Group 6) compared to *L. crispatus* (group 7). (C) volcano plots visualizing downregulated (to the left) and upregulated (to the right) miRNAs differentiating the *G. vaginalis*- and *L. crispatus*-dominated microbiome

**Table 2.** Vaginal dysbiosis-regulated miRNAs*, their validated mRNA targets, and overlap with the HIV Interactome.

| Vaginal microbiome composition affecting miRNA expression | Number of DE miRNAs and their direction of regulation | Number of DE miRNAs with validated targets and their direction of regulation | Number of miRNA-targets and their putative direction of regulation | Number of miRNA-targets overlapping the HIV interactome |
| --- | --- | --- | --- | --- |
| BV vs Normal | 414 up | 141 up | 965 down | 442 |
|  | 174 down | 37 down | 190 up | 50 |
| Intermediate vs Normal | 19 up | 3 up | 32 down | 19 |
|  | 23 down | 5 down | 16 up | 8 |
| BV vs Intermediate | 3 up | 0 up | 0 down | 0 |
|  | 21 down | 6 down | 108 up | 75 |
| GV-dominated vs LC-dominated | 10 up | 3 up | 57 down | 33 |
|  | 14 down | 7 down | 120 up | 79 |

BV= bacterial vaginosis, GV= *Gardnerella vaginalis*, LC= *Lactobacillus crispatus* and DE= differentially expressed. \*FDR <0.01.

**Table 3.** Vaginal dysbiosis-driven differential miRNA dysregulation in directions experimentally confirmed to promote HIV infection.

| DE miRNA | False discovery rate, fold change |  |  | Expected effect on HIV risk based on direction of miRNA dysregulation (46) |
| --- | --- | --- | --- | --- |
|  | BV vs normal | BV vs Intermediate | GV vs LC* |  |
| miR-198↓ | <0.0001, -14.63 | <0.01, -3.12 | <0.001, -10.52 | ↑ HIV replication in monocytes |
| miR-155-5p↑ | <0.05, 1.99 | NS | NS | ↑ reactivation of latent HIV-1 |
| miR-1236-5p↓ | <0.0001, -2.87 | <0.05, -1.74 | NS | ↑ HIV replication in monocytes |
| let-7c↑ | <0.05, 1.70 | NS | NS | ↑ HIV replication in T cells |
| * *G. vaginalis* -dominated vs. *L. crispatus* -dominated metagenome |  |  |  |  |

The BV versus normal Nugent score comparison revealed 141 upregulated EV-miRNAs targeted 965 genes, while 37 downregulated EV-miRNAs targeted 190 genes. The comparison between intermediate and normal Nugent scores found 3 upregulated and 5 downregulated EV-miRNAs, targeting 32 and 16 genes respectively. In the BV versus intermediate Nugent score comparison, 6 downregulated EV-miRNAs were found to target 108 protein-coding genes, with no upregulated EV-miRNAs identified. When stratified by *G. vaginalis*-dominated versus *L. crispatus*-dominated metagenomes, the comparison showed 10 upregulated miRNA (8 overlapped with BV by Nugent score) and 14 downregulated miRNA (all 14 overlapping with the BV by Nugent score). Overall, 92% of the *G. vaginalis* dominance dysregulated miRNAs overlapped with the BV by Nugent score dysregulated genes. Among the those 3 up- and 7 downregulated EV-miRNA had targets validated by Luciferase reporter assay, including targeting 57 and 120 genes respectively. Importantly, the significant down- or upregulation of four EV-miRNAs by BV or by BV and *G. vaginalis* dominance alike, could predict direct facilitation of HIV infection, including enhancing viral replication, monocyte permissiveness to productive HIV infection and reactivation of latent HIV, based on published experimental evidence (**Table 3**).

No miRNAs were differentially expressed with FDR <0.01 when comparing visits that followed metronidazole treatment stratified by normalized versus persistent BV Nugent score.

### 3.2. EV-miRNAs from the Intermediate and BV Nugent score categories target functional networks of macrophage chemotaxis and T cell biology, respectively

The BV versus normal Nugent score comparison of DE EV-miRNA targets identified 34 GOIS terms classified into 8 functional networks (**Fig. 3A**). The two most significant networks identified by ClueGo analysis were associated with ‘lymphocyte differentiation’ (p-value corrected = 3.16 x 10^-11^) and ‘T cell activation’ (p-value corrected = 1.03 x 10^-11^) (**Fig. 3B**). From the intermediate versus normal comparison, a total of three downregulated genes (CSF1, MMP2, MST1) were related with 2 terms grouped together in functional network ‘Chemotaxis’. The BV versus intermediate Nugent score comparison revealed miRNA targets enriched in the functional groups "T cell co-stimulation" (encompassing two terms) and "T-helper 17 cell differentiation" (encompassing one term). The modular enrichment analysis (**Fig. 4A**) and ClueGo analysis (**Fig. 4B**) of validated gene targets identified 6 functional groups from the *G. vaginalis*-*L. crispatus* comparison, including Th17 differential and T cell migration with enrichment for gene with common direction as in the BV-enriched pathways (ACVR2A, ATM, FOXO3, IL6, LIF, MAPK14, NFKBIZ). The overlp between the BV-enriched and G. vaginalis enriched genes is shown in **Table 4**.

**Figure 3:**
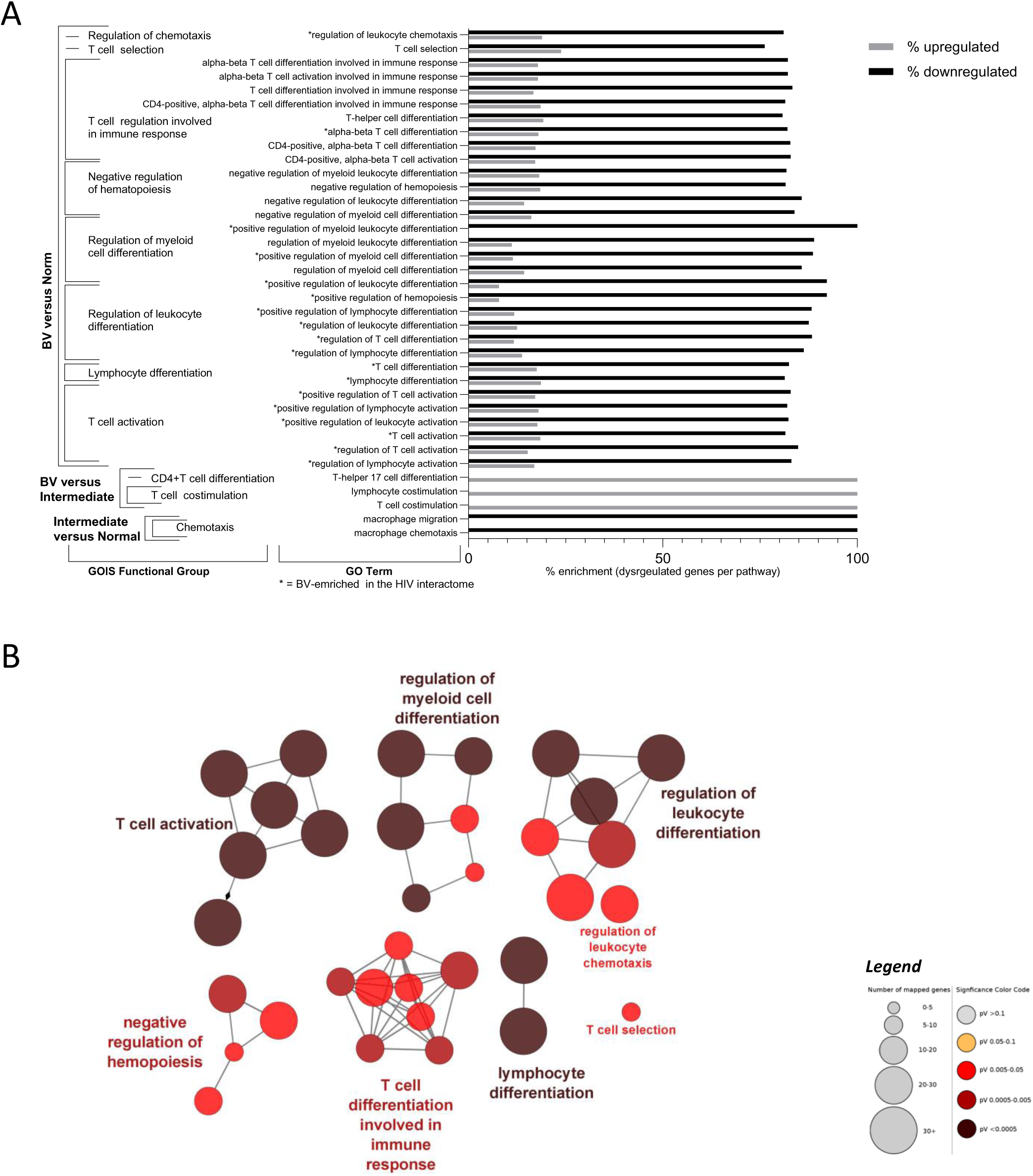
Pathway analysis of DE miRNA targets and enrichment with HIV interactome genes: (A) Bars represent the enrichment by downregulated and upregulated genes (% dysregulated genes per pathway) for each of the GOIS Functional groups and GO terms associated with DE EV-miRNAs in the three comparisons (BV versus Normal, BV versus Intermediate, and Intermediate versus Normal Nugent score. Asterisks denote BV-enriched terms in the HIV interactome. (B) ClueGO functional network analysis of all target genes of DE miRNAs in BV versus normal Nugent score, using the Gene Ontology Immune System (GOIS) database. Nodes represent terms associated with each of the eight functional networks; bubble size indicates the number of genes per term (ranging from 5 to 30+), and color represents significance as shown in the legend insert.

**Figure 4.**
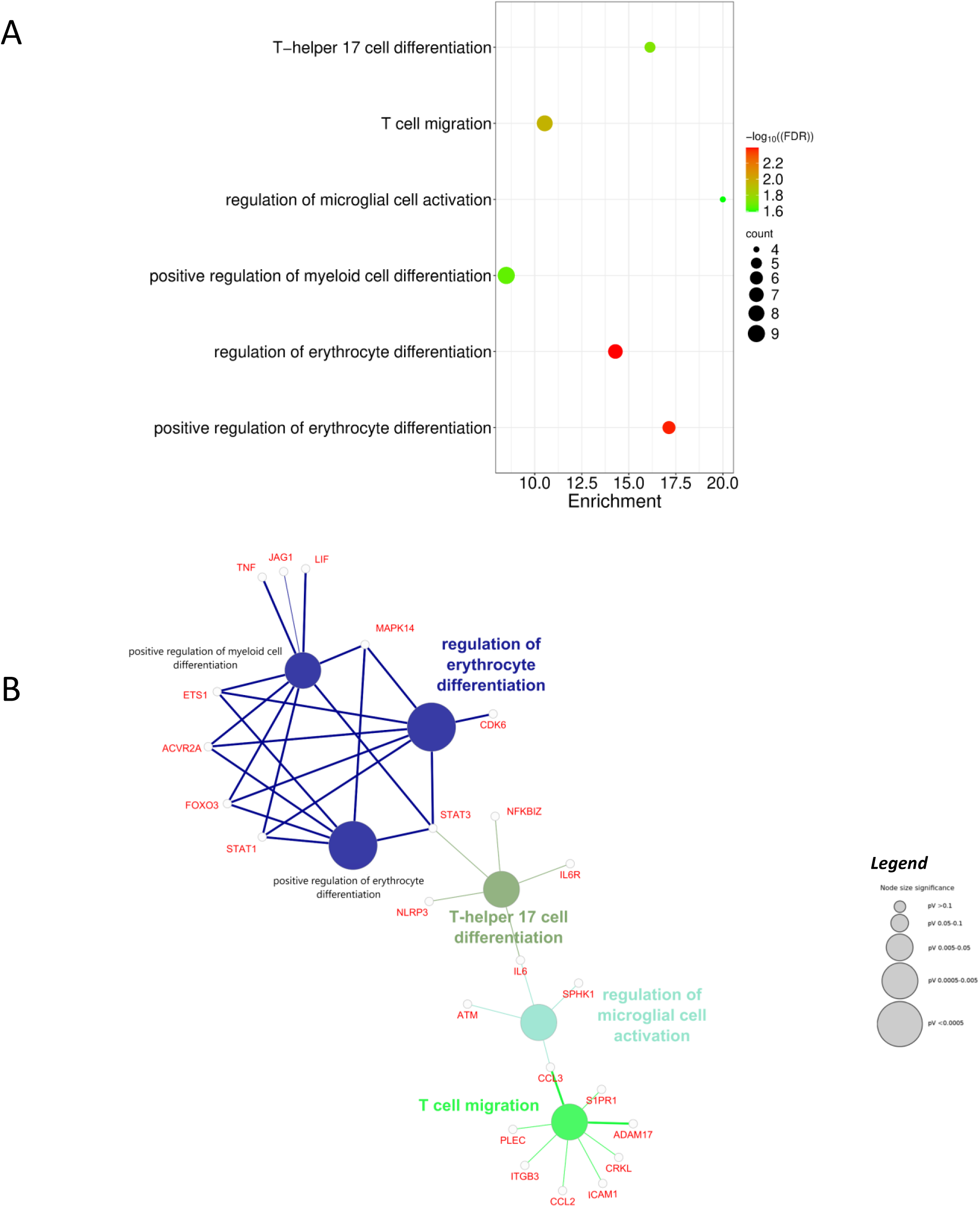
(A) Modular Enrichment Analysis (MEA) of validated targets of DE EV miRNAs in the GV-dominated vs. LC-dominated metagenome comparison. The bubble plot shows enriched GOIS terms (Y-axis) and percentage of genes per term (X-axis). Bubble color reflects significance (–log₁₀ FDR); size indicates number of genes per term. (B) ClueGO functional network of validated targets from (A) using the GOIS database. Colored nodes represent functionally grouped terms; labeled genes contribute to enrichment. Node size reflects significance (p > 0.1 to p < 0.0005); thicker edges indicate stronger relationship,

**Table 4.** Validated DE-miRNA Targets Shared and Unique to BV vs. Normal and GV vs. LC, Contributing to GOIS Enrichment.

|  |  |  |
| --- | --- | --- |
| Sharing Status | Group | Gene Symbol |
| *shared between groups* | [GV protein down] & [BV protein down] | ACVR2A, ATM, FOXO3, IL6, LIF |
|  | [GV protein up] & [BV protein up] | MAPK14, NFKBIZ |
|  | [GV protein up] & [BV protein down] | ADAM17, TNF |
|  | [GV protein up] & [BV up & down] | CCL2, CDK6, ETS1, IL6R, JAG1, STAT3 |
|  | [BV protein up] & [GV protein down] | ICAM1 |
| *unique to each group* | [GV protein up] | CRKL, ITGB3, PLEC, SPHK1 |
|  | [BV protein up] | AHR, CEBPA, CX3CR1, EFNB1, GREM1, KLF6, PADI2, SMARCC1, SOS1, XRCC6, YY1, ZAP70 |
|  | [GV protein down] | CCL3, NLRP3, S1PR1, STAT1 |
|  | [BV protein down] | ABL2, ACVR1B, ADAM10, ADORA2A, ANXA1, AP3B1, ATG5, AXL, BAX, BCL10, BCL6, BTK, CADM1, CARD11, CASP3, CASP8, CCL1, CCND3, CCR7, CD226, CD40, CD44, CD47, CDKN1A, CREB1, CSF1, CTNNB1, CTSC, CUL4A, CXCL12, CXCL8, CYP19A1, DLG1, DLG5, DLL1, DUSP10, EDN1, EFNB2, EOMES, FBN1, FBXW7, FOS, FOXN1, FOXP1, FYN, FZD5, GDNF, GLI2, GPR137B, HDAC4, HIF1A, HLA-G, HMGB1, HMGB2, HOXA5, HOXA7, HOXA9, HOXB8, ID2, IFNG, IGF1, IGF2, IGFBP2, IKZF1, IL10, IL12B, IL1A, IL1B, IRF1, ITCH, JAK1, JUNB, KIF20B, LEF1, MAFB, MALT1, MAP3K8, MAPK1, MAPK11, MEF2C, MEIS1, MERTK, METTL3, MIF, MITF, MLH1, MMP8, MST1, NCOR2, NCSTN, NFIL3, NFKBIA, NOTCH2, PBRM1, PCK1, PHF10, PIAS3, PIK3CD, PIK3CG, PIK3R1, PIK3R2, PKNOX1, POU2F2, POU4F2, PRDM1, PTK2, PTK2B, PTPN11, RAC1, RARA, RASSF5, RB1, RCOR1, RHOA, RORA, RUNX1, RUNX2, RUNX3, S100A9, SCRIB, SDC4, SERPINE1, SIRPA, SLC7A1, SMAD7, SMARCA2, SMARCB1, SOCS5, SOCS6, SOD1, SOX4, SPI1, SRC, SRF, ST3GAL1, STAT6, STK11, STK4, TCF3, TGFBR2, THBS1, TIMP1, TLR3, TLR4, TMEM64, TNFAIP3, TNFRSF11A, TNFSF11, TP53, TPD52, TRAF6, TRIB1, TSC1, VCAM1, VEGFC, VSIR, WNK1, WNT10B, WNT5A, XBP1, YES1, ZC3H12A, ZEB1, ZFPM1 |
|  | [BV up & down] | AKT1, BCL2, BMI1, CAV1, CD151, CD274, EGR1, ERBB2, EZH2, HES1, ITGB1, KIT, LAMP1, MTOR, MYB, MYC, NF1, NOD2, PIK3CA, PIK3R3, SFRP1, SMAD3, SNAI2, VEGFA, WNT1, WNT3A |

More detailed information on the targets of DEmiR and associated enriched terms are in the **Supplemental Table 4**.

### 3.3. The targets of EV-miRNAs dysregulated in BV and Intermediate phenotypes overlap with HIV interactome genes and are associated with KEGG terms in infection, leukocyte biology and endometrial cancer

The HIV interactome shares 442 genes predicted to be downregulated and 50 genes predicted to be upregulated as targets of BV-dysregulated EV-miRNAs (see **Supplemental Table 5).** KEGG pathway analysis revealed 172 enriched terms (adjusted p-value < 0.05), organized into 90 functionally associated networks. Notably, the top 30 enriched pathways emphasized bacterial, protozoan, and viral infection processes, including HIV. Additionally, GOIS analysis of these BV-HIV overlapping genes identified 44 enriched terms (adjusted p-value < 0.05), categorized into the eight functionally grouped networks, enriched by BV, including processes related to leukocyte differentiation and activation (**Fig. 3A**). Of note, the GOIS and KEGG enriched terms were predominantly associated with the predicted downregulated genes (see **Supplemental Table 6**).

The intersection of genes between the targets of Intermediate-dysregulated EV-miRNAs and the HIV interactome encompassed 8 and 19 genes predicted to be, up- and downregulated respectively (see **Supplemental Table 5**). KEGG enrichment analysis identified 10 enriched terms (adjusted p-value < 0.001) distributed into 5 functionally associated networks. Enriched terms relevant to the female genital tract health includes terms on ‘adherens junction’, ‘prolactin signaling pathway’, and ‘endometrial cancer’ (see **Supplemental Table 6**). The GOIS analysis did not return data.

### 3.4. The targets of EV-miRNAs dysregulated in G. vaginalis-dominated metagenome impact T cell biology and cytokine signaling

The validated targets of EV-miRNAs dysregulated in GV-dominated microbiome compared to LC-dominated microbiome showed a total of 6 enriched GOIS terms, including 2 terms linked to T cell function, encompassing a total of 24 interrelated genes (**Fig. 4**, **Supplemental Table 7**). In addition, 79 validated targets predicted to be upregulated and 33 predicted to be downregulated by these EV-miRNAs dysregulated in the GV-dominated microbiome overlap with the HIV interactome (**Supplemental Table 5**). These shared targets are enriched in GOIS terms, including those linked to T cell biology, while the KEGG terms highlights several terms associated to FAN related to viral infection, TNF signaling pathway and T17 cell differentiation (**Supplemental Table 6**).

### 3.5. Correlation of DE miRNAs with innate immunity mediators implicated in HIV risk

The results of testing the association of the continuous miRNA levels and cytokines treated as categorical are shown in **Supplemental Table 8.** The top 50 DE miRNAs per cytokine are shown in **Fig. 5A-E**.

**Figure 5.**
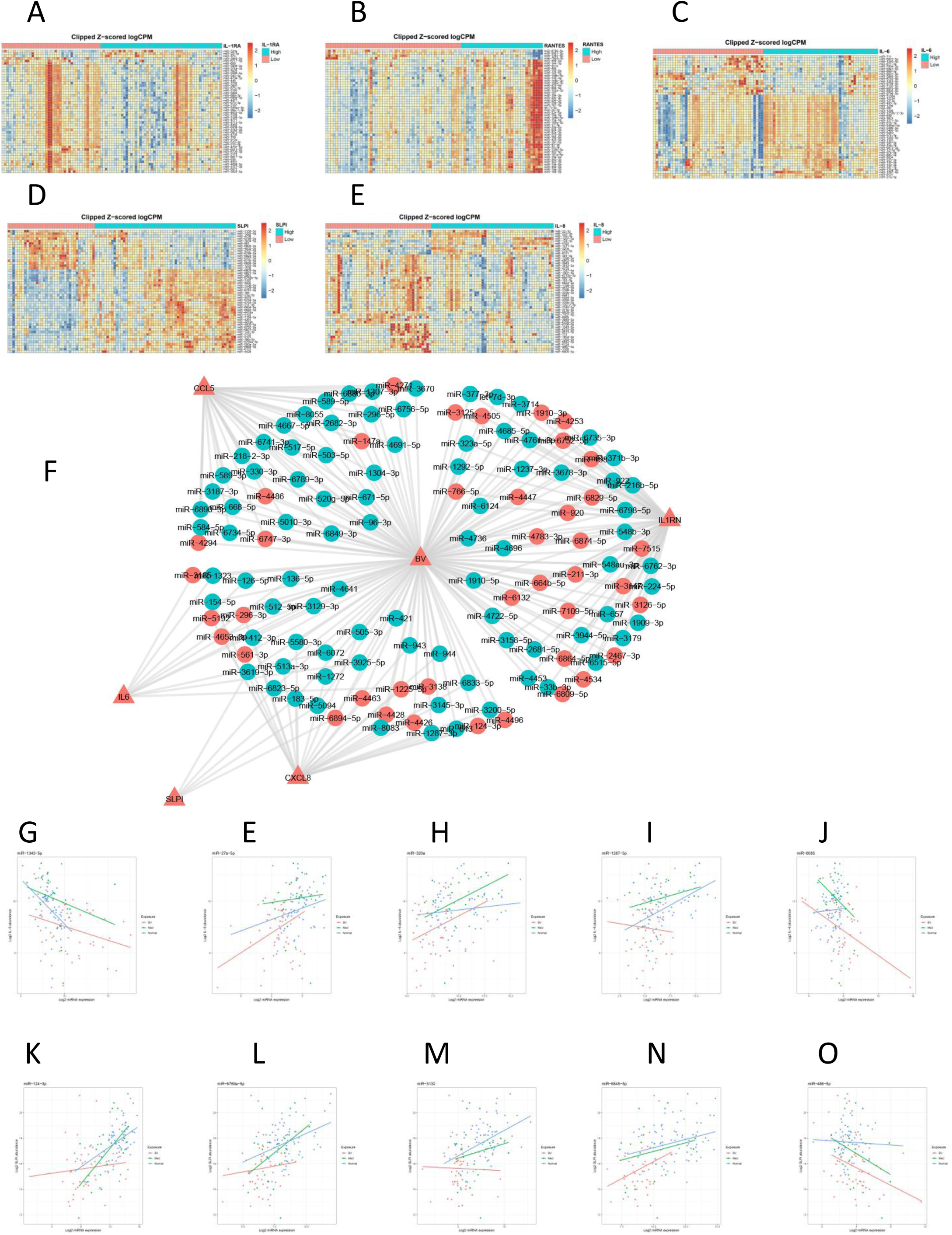
miRNA correlation with cervical innate immunity mediators implicated in HIV risk: (A-D) Heat maps illustrate the top 50 miRNA ranked by adjusted p value of most significant correlation with the protein levels of SLPI (A), RANTES (B), IL-6 (C) and IL-8 (D) and IL-1RA (E). (F) BV-miRNA-cytokine network. The circle symbols represent miRNAs that are correlated with the cytokines connected to them in a network node and that are downregulated (red circles) or upregulated (teal green circles) in women with BV. (G-O) Plots visualizing results from the Hitmna analysis depicting miRNA mediators consistent with BV suppressing IL-8 (G-J) and SLPI (K-O).

The cytokine correlation network shown in **Fig 5F** visualizes the top miRNA and the corresponding cytokines related to Nugent scores based on the combined FDR < 0.05. The network analysis selected the candidate BV -> mir -> cytokine connections according to the following criteria: (1) the microRNA is affected by BV in comparison with normal Nugent Score (FDR < 1%) and (2) the cytokine targeted by the miRNA.

The Spearman correlation between the Nugent categories showed significant negative association (p value and FDR<0.05) between BV and SLPI (rho=-0.5342525) and IL-8 (rho=-0.2758373) and therefore they were selected for the Hitman mediation analysis including all miRNAs (**Supplemental Table 2**). The top 5 miRNAs mediating the downregulation of the cytokines by BV are shown in **Fig. 5H-J** for IL-8 (Mir 1343-5p, miR-27a-5p, miR-320-a, miR-1287-5p, mir-8085) and **Fig. 5K-O** for SLPI (miR-124-3p, miR-6769a-5p,miR-3132, miR-6845-5p, miR-486-5p).

## 4 Discussion

This study provides robust clinical evidence for the role of vaginal bacterial dysbiosis in modifying exosome miRNAs with impact on immunity and susceptibility to HIV. Cervicovaginal bacteria differentially alter miRNA expression in ectocervical cells *in vitro* (36) and significant changes in the miRNA cargo of exosomes were observed in an experimental human model of vaginal colonization with healthy lactobacilli versus BV pathogens (2). These early studies provided proof for the causative role of bacteria in altering the cervicovaginal miRNAome, that needed clinical validation. Prior studies of cervicovaginal miRNAs suffered from smaller size and limited dimensions. One study investigated expression of only two miRNAs in cervicovaginal lavage (CVL) from 45 women with cervical cancer (37). Another study investigated the presence of exosomes in vaginal fluid and showed vaginal exosomes have an effect on HIV *in vitro* but did not investigate the miRNA content of the exosomes (38). A study on BV and pre-term birth, attempted to investigate expression of 800 miRNA in cervical cells from 80 women (39). Using Nanostring nCounter methodology for miRNA detection, only 74 miRNA probes were detectable in at least 60% of all samples(39). One miRNA (miR-494) was associated with a pre-calculated bacterial index and 27 miRNAs – with a cytokine index (p<0.05 and FDR<0.01) (39).

In contrast to all prior studies limited by size and scope, the evidence presented here is based on a global transcriptome analysis detecting 2081 human miRNAs coupled with conservative approach to target validation, robust clinical diagnosis, and deep-sequencing confirmation for the role of *G. vaginalis*, which is considered a signature BV pathogen (1). This allowed us to identify the magnitude and direction of over 500 dysbiosis-regulated mRNAs along with over 1000 gene targets, their putative direction of dysregulation, over half overlapping with the HIV interactome. Unlike other studies we also correlated the global miRNAome with the expression levels of 5 proteins (SLPI, RANTES, IL-8, BD2 and IL-1RA, previously identified as predictors on HIV acquisition risk(28, 29, 40) in one of the largest longitudinal cohorts that followed HIV negative women over a median period of 21.5 months (28, 40-42). We applied a novel Hitman analysis to identify the top 5 miRNAs mediators consistent with the immunosuppressive effects of BV on SLPI and IL-8 levels in the same samples that were used for miRNA analysis. SLPI is an important mediator of innate immunity with anti-inflammatory properties and its levels are decreased in any sexually transmitted infection and BV(43, 44). We add mechanistic evidence for the causative link with BV. IL-8 is a chemokine essential for neutrophil migration to the site of infection.(45) While epithelial cells react to individual BV and sexually transmitted pathogens with increased production of IL-8, the dampening of this first line response by BV through increased miRNA production offers a mechanistic path for evading the host immunity and preventing pathogen clearance, allowing for long-term BV bacteria colonization.

Our study supports the hypothesis that EV-miRNAs differentially expressed in BV and in *G. vaginalis* dominated microbiomes may represent a molecular mechanism for the strong epidemiologic association of BV with HIV infection (2). In this study the global transcriptome analysis revealed that more than a quarter of all known human miRNAs are dysregulated under the condition of BV and at least a log difference between the number of miRNAs affected by BV compared to those affected by the intermediate microflora conditions (Nugent score 4-6). Emerging evidence confirms the role of specific miRNAs in HIV infection (46). A review summarized the experimental evidence of direct effect of 23 miRNAs reported to target host-dependence factors and modulate HIV replication either negatively or positively(46). Four of these miRNAs were DE in BV and BV along *G. vaginalis* dominated and /or intermediate microflora, leading to down or upregulation of genes consistent with direct enhancement of HIV replication, latent virus reactivation or suppressing monocyte infection.

Among those mir-198 requires a special attention, because it exemplifies an over-a-log fold downregulation by both Nugent- and metagenome-defined dysbiotic conditions. This miRNA has been experimentally proven to impair HIV infection of monocytes (46), and thus, its reduction by BV (and *G. vaginalis)* may represent another underlying mechanism of BV-viral synergisms, evading the normally low permissiveness of monocytes to HIV infection. The vast overlap with the HIV interactome leading to enrichment of key pathways of T cell and monocyte immunity deserves further studies to identify prevention mechanisms.

## 5. Conclusion

The vaginal microbiota plays a role in defining the miRNA cargo of mucosa-derived extracellular vesicles that can exert epigenetic control over protein expression. miRNAs dysregulated by dysbiotic conditions diagnosed by abnormal Nugent score or metagenomic dominance of bacterial species characteristic of bacterial vaginosis may underlie the HIV risk. These findings open the door to new strategies for prevention of BV-associated complications and morbidities, including HIV preventions.

## Author Contributions

RNF conceived the study, obtained funding and drafted the manuscript. GFD designed and led the clinical trials that provided the biospecimens and metadata for this study. PFTCM completed pathway analyses, functional and network analysis and contributed methods and graphs for figures 3 and 4. JD and PH contributed bioinformatics and statistical analysis in R and contributed graphs to figures 1, 2, and 5. HY and SN contributed to the miRNA data acquisition. XG contributed biostatistical expertise. CM and PLC contributed epidemiologic expertise. RS and RB contributed clinical expertise. All authors provided editing and have read and approved the submission of this article. The authors would like to pay tribute to the late Dr. Robert Salata, Case Western reserve University, Ohio, USA, a devoted infectious disease clinician and internationally renowned scholar, who consulted on the early study design and tragically passed away before the study was completed.

